# Performance Of Children With Autism In Parent-Administered Cognitive And Language Exercises

**DOI:** 10.1101/146449

**Authors:** Rita Dunn, Jonah Elgart, Lisa Lokshina, Alexander Faisman, Edward Khokhlovich, Yuriy Gankin, Andrey Vyshedskiy

**Author notes:** **Corresponding author:** Andrey Vyshedskiy, Ph.D., Boston University, Boston, USA.

## Abstract

There is a broad scientific consensus that early and intensive therapy has the greatest chance of positive impact on an individual with autism spectrum disorder (ASD). However, the availability, quality, and general funding for early intervention programs is often lacking, leaving newly diagnosed children without adequate and sufficient therapy during the most critical early period of their development. Parent-administered iPad-assisted therapy has the potential to reduce the gap between the amount of therapy recommended for children with ASD and the amount they receive. However it is unclear how ASD severity and age influence a child’s ability to engage with and learn from computerized cognitive exercises. In this manuscript, we describe data from a tablet-based therapeutic application administered by parents to 1,514 young children with ASD over the course of four to twelve months. We report that older children and children with milder forms of ASD performed better and progressed faster in cognitive and language exercises. However, most children were able to engage with and learn from exercises independent of their age or ASD severity. This data confirm that tablet-based cognitive and language exercises can be successfully administered by parents to children as young as two years of age over the course of many months independent of ASD severity.

## Introduction

The Centers for Disease Control estimates that 1 in 68 (Wingate et al., 2014), children is affected by autism spectrum disorder (ASD), a neurological disorder that disrupts early development in cognition and communication. Approximately two-thirds of children with ASD grow up to have a significant cognitive and social impairment, and difficulty in acquiring new adaptive behaviors (Yeargin-Allsopp et al., 2003). There is broad scientific consensus that early and intensive behavioral intervention can result in sizeable gains in cognitive, communication, social, academic, and adaptive skills, and has the greatest chance of significantly improving outcomes, sometimes even resulting in a complete loss of diagnosis (Eldevik et al., 2010; Peters-Scheffer, Didden, Korzilius, & Sturmey, 2011; Virués-Ortega, 2010). Accordingly, the American Academy of Pediatrics (AAP) recommends universal screening of 18- and 24-month-old children for ASD, and also that individuals diagnosed with ASD begin to receive no less than 25 hours per week of treatment within 60 days of identification (Maglione et al., 2012). Despite the AAP recommendation, two-thirds of US children on the autism spectrum under the age of 8 fail to get even the minimum recommended treatment (Hume, Bellini, & Pratt, 2005) because of major problems with the availability, quality, and general funding for early intervention programs (Bibby, Eikeseth, Martin, Mudford, & Reeves, 2002; Jacobson, 2000; Johnson & Hastings, 2002). Since the AAP’s 2007 recommendation of universal, early screening, there has been a sharp increase in demand for ASD-related services (58% on average, Ref. (Wise, Little, Holliman, Wise, & Wang, 2010). However, according to a recent study, most states have reported an enormous shortage of ASD-trained personnel, including behavioral therapists (89%), speech-language pathologists (82%), and occupational therapists (79%) (Wise et al., 2010). In many states children are getting less than 5 hours per week of service, and this immense shortage disproportionally affects African American and Latino children (Wise et al., 2010). Families of newly diagnosed children often face lengthy waitlists for therapy, leaving children without treatment during the most critical, early period of development. The failure to provide adequate early intervention services ends up costing society (USA) an estimated $126 billion per year (Mandell & Knapp, 2012).

The use of technology in the delivery of much needed therapy promises to be effective for shrinking the gap between the therapy that is recommended for children with ASD and the amount they receive. Recently, there has been a trend towards computer-based interventions which capitalize on the often-observed preference that children with ASD show for flat screen information (Bernard-Opitz, Sriram, & Nakhoda-Sapuan, 2001; Schreibman, Whalen, & Stahmer, 2000; Shukla-Mehta, Miller, & Callahan, 2010) as well as evidence suggesting that tablets specifically have been a great learning tool for children on the spectrum (Jowett, Moore, & Anderson, 2012; Kagohara, Sigafoos, Achmadi, O’Reilly, & Lancioni, 2012). However, since there is scant research on computer-based interventions for children with ASD, many questions remain to be answered: at what age are children diagnosed with ASD capable of being fully engaged with computer-based interventions? Critically, can the neediest population of newly diagnosed children as young as 2 with severe ASD symptoms engage with and benefit from such an intervention? Does the severity of ASD symptoms influence engagement?

In order to answer these questions, we developed a tablet-based therapeutic application for children with ASD called *Mental Imagery Therapy for Autism* (MITA) (Vyshedskiy & Dunn, 2015). In this report, we describe data from a feasibility study of MITA that included 1,514 children with ASD who worked with the application for four to twelve months. We discuss MITA usability, feasibility, fidelity of implementation and promise of outcomes. To our knowledge, this report is the first to study tablet-based cognitive exercises administered to 2- year-old toddlers with autism and it is also the largest study of an early parent-administered intervention tool to toddlers with autism.

MITA’s tasks and overall curriculum are grounded in some of the best scientifically supported and established, evidence-based therapies for ASD (Wilczynski et al., 2009), drawing most heavily from Applied Behavioral Analysis (ABA) and Pivotal Response Training (PRT). MITA focuses on one of the four key, or “pivotal,” areas of development targeted by PRT that in turn affects a wide range of behaviors. MITA’s educational objective is to develop a child’s ability to notice and respond to multiple cues presented simultaneously. To understand this ability, imagine that you are instructed to *pick up a red crayon that is under the table.* This may seem like a trivial task, but in order to accomplish it successfully, you need to notice three different features, or “cues,” of the object: its color (*red*), its shape (*crayon*) and its location (*under the table*). You must then mentally integrate all three pieces of information into a new mental image, *a red crayon under the table,* in order to take the correct action. The ability to integrate multiple cues is highly developed in individuals not afflicted by ASD well before the age of 6, but it is known to be a common challenge for children on the spectrum (Lovaas, Schreibman, Koegel, & Rehm, 1971). As a consequence, ASD symptoms often include a phenomenon called stimulus overselectivity, whereby an individual focuses on only one aspect of an object or environment while ignoring others (Lovaas, Koegel, & Schreibman, 1979; Ploog, 2010; Schreibman, 1988). When asked to *pick up a red crayon under the table*, a child with ASD may hyper-attend to the cue “crayon” and ignore both its location and the fact that it should also be red, therefore picking up any available crayon. It is often said that individuals with ASD “can't see the forest for the trees”: they pay too much attention to specific parts, get lost in the details and miss the whole picture (or Gestalt). The consequences of attempting to navigate the world with an impaired ability to respond to multiple cues can be profound and can affect virtually every area of functioning (Lovaas et al., 1971). However, using PRT to develop responsivity to multiple cues has been shown to *reduce* stimulus overselectivity and, most importantly, to lead to improvements in general learning (Burke & Cerniglia, 1990; Wingate et al., 2014). In addition to developing a child’s ability to respond to multiple cues, MITA also aims to train receptive language, starting with simple vocabulary, and progressing towards higher forms of language, such as adjectives, verbs, pronouns, and syntax.

## Methods

We present data from an observational study of a tablet-based therapeutic application called *Mental Imagery Therapy for Autism* (MITA) administered by parents to 1,514 young children with ASD over the course of four to twelve months. MITA was developed by ImagiRation from 2013 to 2016 and made available for free at all major app stores in February of 2016. In the first year, MITA was downloaded 70,325 times, indicating a significant interest in supplemental therapy for ASD.

Once MITA was downloaded, the user was asked to register and provide demographic details, including the child’s diagnosis as well as month and year of birth. During twelve months (from February of 2016 to February of 2017) MITA was registered 41,690 times (59% of downloads). Table 1 shows the distribution of registration over operating systems and device types.

**Table 1.** MITA download statistics

| Table 1 |  |
| --- | --- |
| *MITA download statistics* |  |
| Device or platform | Percent downloads |
| Android phones | 35% |
| Android tablets 10 inches+ | 21% |
| Android tablets 7 inches to 10 inches | 18% |
| iPad tablet | 16% |
| iPhone | 5% |
| Amazon tablets | 5% |
| All downloads from Google Play | 74% |
| All downloads from Apple app store | 21% |
| All downloads from Amazon app store | 5% |
| All tablet devices | 60% |
| All smartphones (including larger smartphones) | 40% |

### Subjects

From the pool of potential study subjects who downloaded MITA, we selected subjects based on the following criteria:

1) **The subject’s parent must have filled out at least two questionnaires at least three months apart.** Since our application was available for free to the general public, we expected a large volume of downloads by people of widely ranging commitment. We needed a benchmark to discern those who had serious intentions in working with a therapeutic application and those who did not. Approximately one month after the first use of MITA and no sooner than 100 puzzles had been solved, parents were required to complete the 77-question Autism Treatment Evaluation Checklist (ATEC, (Rimland & Edelson, 1999). Subsequently, parents were asked to complete ATEC at three month intervals in order to continue their use of MITA. Parents who invested the time to faithfully fill-out an extensive ATEC questionnaire on two separate occasions demonstrated such minimal commitment. This automatically selected subjects who used MITA for at least four months. As of February 2017, out of the 41,690 registered users, 30,248 had already used MITA for at least 4 months and therefore could potentially have completed two evaluations. The number of subjects who actually completed two or more evaluations was 2,525 (8% of all potential subjects).
2) **The subject’s parent must have self-reported the diagnosis as ASD.** Since our primary interest is early intervention for ASD, only data from ASD subjects were analyzed for this report. Out of the 2,525 parents who completed at least two questionnaires, 1,839 (73%) self-reported their child’s diagnosis as ASD. Other subjects reported diagnoses of various other neurodevelopmental disorders or that they were not yet diagnosed. Some subjects chose not to report a diagnosis since this was not a required field.
3) **The subject must have been 12 years of age or younger at the time of registration.** Since we are interested in the effects of early intervention, we decided to limit our analysis to subjects who were 12 years of age or younger at the time of the first questionnaire. Therefore, we excluded another 323 subjects because of age. Thus, the total number of subjects included for analysis was reduced to 1,514 (5% of all potential subjects).

The number of subjects in each age group is indicated in Table 2. Seventy nine percent of subjects were male.

**Table 2.** The number of subjects in each age group

| Table 2 |  |  |  |
| --- | --- | --- | --- |
| *The number of subjects in each age group* |  |  |  |
| Age | Age marked on the graphs | Number of subjects in each age group | % subjects in each age group |
| 1.5 to 3.9 | 2 to 3 | 787 | 52 |
| 4 to 5.9 | 4 to 5 | 453 | 30 |
| 6 to 12 | 6 to 12 | 274 | 18 |
| Total |  | 1514 | 100 |

### Experimental interventions

All subjects in this study used a tablet-based therapeutic application called *Mental Imagery Therapy for Autism* (MITA) developed by ImagiRation. MITA consists of nine different developmental activities, described in detail in Supplemental Materials. All of the activities follow a systematic approach to train the skill of multiple cue responding, and eight of the activities provide this training outside of the verbal domain (Vyshedskiy & Dunn, 2015). This unique feature allows the activities to be within reach of those individuals who are either nonverbal or minimally verbal. While these children may not be able to follow an explicit verbal command (such as “*pick up the red crayon under the table*”), results from our studies have demonstrated that they can follow a command implicit in the visual set-up of the puzzle.

To teach children to follow implicit visual commands that require attending to multiple cues, the MITA program starts with puzzles that require attending to one cue, such as color (Fig. 1A) or shape (Fig. 1B). Once a child shows adequate proficiency in attending to a variety of single cues, MITA activities progress in difficulty by requiring attention to two cues simultaneously, such as both color and shape (Fig. 1C) and eventually to three or more cues.

**Figure 1.**
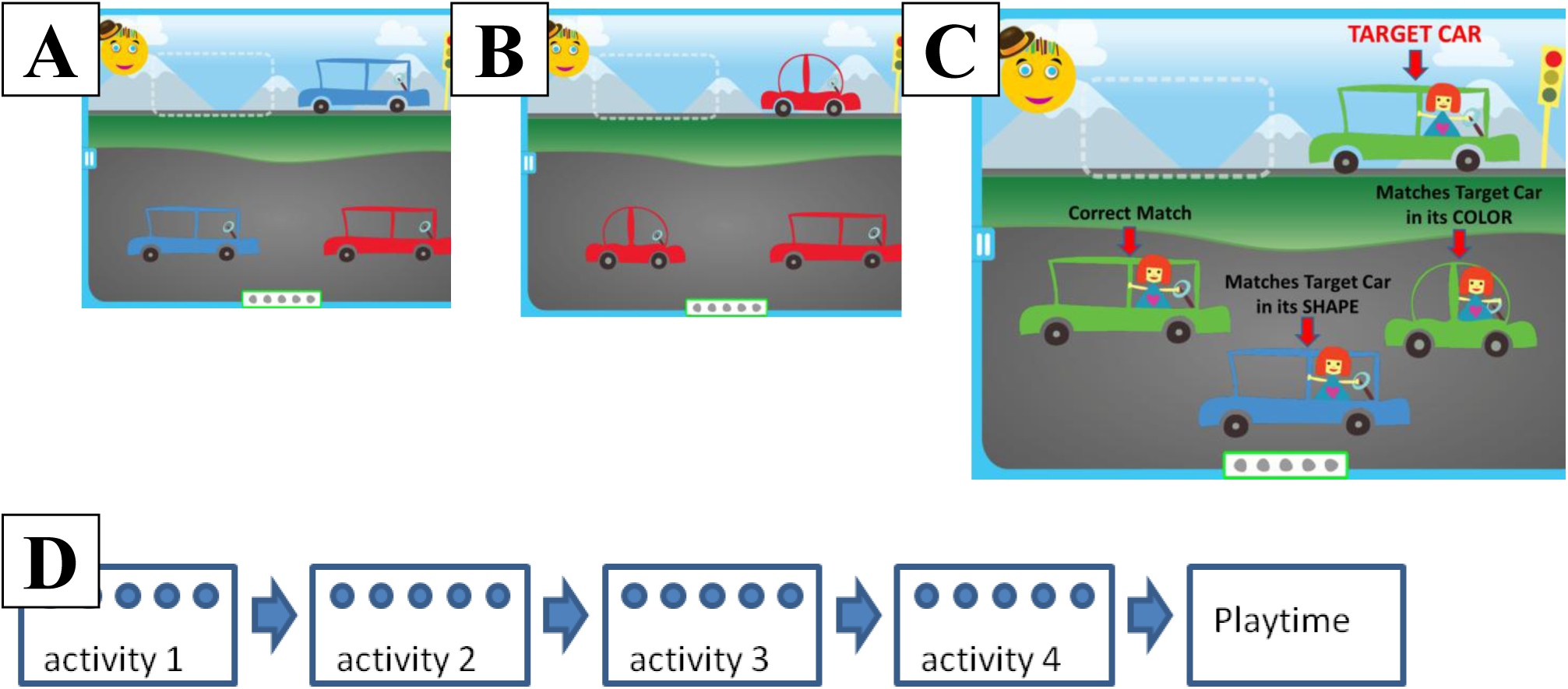
Examples of puzzles with increasing complexity. (a) A child must notice one cue (color) in order to correctly match the car at the top of the screen. (b) A child must notice one cue (shape) in order to correctly match the car. (c) A child must notice two cues (color and shape) in order to correctly match the car (note: the overlaid text and arrows are not seen by the child). Notice that hyper-attending to a single cue (only the shape or only the color of the vehicle) and ignoring the other cues is insufficient for finding the correct match. (d) A default daily session consists of 4 activities followed by Playtime. Each activity contains 5 puzzles marked by a circle on the diagram. See supplemental materials for the description of each activity.

In addition to color, shape and size, MITA activities sometimes require attention to an object’s orientation in space, the number of objects, as well as minor visual details. The choice of these particular stimuli was made to reflect those commonly used by PRT behavioral therapists who intentionally structure the therapeutic environment to include objects of various color, shape and size and then ask the child to find an object based on two (or more) of these features (Hiniker, Daniels, & Williamson, 2013).

Each of the nine MITA activities consists of multiple levels, starting with easier levels that require attending to a single cue, moving on to intermediate levels that require attending to two cues and culminating in challenging levels that require attending to three or four cues at a time. Most activities have as many as 50 levels which range from easy to difficult in a gradual and systematic manner. Please refer to the Supplemental Materials for a detailed discussion and a variety of examples of MITA activities and levels.

Progression from one level to the next is adaptive. A child must demonstrate a high level of proficiency before being advanced to the next level by the adaptive algorithm. Progression through levels is not always upward; a child who is failing at a particular level could be demoted down to the previous level. A child who is showing only moderate achievement may stay at a single level until mastery is demonstrated. The gradual increase in difficulty coupled with a strict assessment of mastery before progressing to the next step is ideal for training multiple cue responding.

#### Daily session

MITA activities are organized into daily sessions, Fig. 1D. The minimum length of the daily session is determined by the number of activities presented on the screen as well as the number of puzzles in each activity. The minimum length of the daily session automatically adjusts based on age and performance, with predefined benchmarks based on recommendations from ABA specialists. The initial number of activities in each session was set to four activities for children younger than 6 years of age, and six activities for children older than 6. The number of activities automatically increased by one (until the maximum of six) once the child had shown progress by completing at least half of the difficulty levels in any two activities. The initial number of puzzles within each activity was set to five puzzles. In addition to these automated settings, the minimal number of activities could be manually adjusted to anywhere from one to six activities per session, and the number of puzzles within each activity could be manually adjusted to 5, 10, 15 or 20 puzzles. Once children complete all the puzzles in each of the daily session’s activities, they are rewarded with Playtime. Note that the maximum number of puzzles per session is not limited: children can reenter any activity and continue working with puzzles for as long as they want.

Following the recommendations of consulting ABA specialists, we advised parents to exercise with MITA on a daily basis for approximately 10 minutes each day until the child is able to reach the top difficulty level in each activity, which is likely to take as long as two to three years.

### Outcome Measurements

We were interested in looking at how subjects performed at their best, as well as their average performance over time. Thus, we created two measurements of performance: 1) The Highest Achievement score and 2) The Average Performance score.

#### 1) The Highest Achievement score

As discussed above, MITA uses adaptive algorithms to advance a subject through the levels over time. Because inconsistent patterns of attention, intellectual function and behavior are a hallmark of children on the autistic spectrum, we expected to see many subjects fluctuate in their level in any particular activity. In fact, our data show exactly such vacillations. We have observed drops of five or more difficulty levels in 342 subjects (23%) and drops in three or more difficulty levels in 732 subjects (49%).These drops were intermittent and unpredictable. For example, a subject who was doing well for months and who had reached and sustained level 20 in a particular activity may then experience a downswing and descend to level 12 over the course of weeks before beginning to rebound. Using the subject’s current level (12) would therefore not be an adequate gauge of that child’s overall achievement in this activity since he had previously succeeded in a more challenging level. To account for possible downswings, the Highest Achievement score was calculated based on the highest level reached (and sustained over the course of at least three daily session) instead of the current level. Specifically, the Highest Achievement score was calculated as a sum of sustained maximum levels reached by the child in each of the nine activities normalized by the maximum possible level in each activity. Reaching the highest possible level of difficulty in all activities corresponds to the Highest Achievement score of 100. A score of 1 corresponds to not progressing beyond the easiest level in any activity.

Since MITA consists of puzzles which incrementally increase in difficulty, it is in some sense analogous to an IQ test, and therefore, the Highest Achievement Score is analogous to an IQ score. Of course, this analogy only works after the subject has worked with MITA long enough for the adaptive algorithm to have sufficient data to match the subject’s actual ability level with the activity’s difficulty level. Furthermore, as the subject ages and his intellectual faculties improve, we would expect his Highest Achievement score/“MITA IQ score” to increase as well.

#### 2) The Average Performance score

Whereas the Highest Achievement score captures a child’s best performance and disregards fluctuations, the Average Performance score encapsulates a child’s overall performance over all daily sessions. Performance is assessed after the completion of every puzzle with a score inversely proportional to the number of errors and normalized by the number of answer choices. For example, in a puzzle with one task (e.g. find the matching animal) and three answer choices (one correct and two decoys), the performance score could be 100% (subject found the correct answer on the first try), 50% (subject found the correct answer on the second try) or 0% (subject found the correct answer only after exhausting all possible options). Making more than three errors in a puzzle with only three answer choices corresponds to a performance score of 0%. Accidental drags and drops did not count as incorrect answers because we did not want to penalize subjects for poor fine motor skills. The performance scores for all the puzzles solved in a daily session are averaged into a daily performance score, which are in turn averaged into the Average Performance score.

Since the Average Performance score is an average of all daily sessions, it can be easily affected by aberrant days when the child is cognitively disengaged or going through a downswing due to sickness or a period of seizures. To avoid factoring in such anomalous activity into the score, we established a minimum threshold of 20 completed puzzles since this number corresponds to our pre-set minimum daily session (4 activities with 5 puzzles in each activity). Days when a subject was unable to solve the minimum of 20 puzzles were excluded from analysis because the subject failed to demonstrate the minimum level of commitment to engaging with the app on that day.

### Autism Severity Measurements

Each subject was assigned their autism severity group mild, moderate, or severe based on their 77-question Autism Treatment Evaluation Checklist (ATEC, (Rimland & Edelson, 1999) score. Parents were asked to complete ATEC at three month intervals in order to continue their use of MITA. In addition, parents were able to complete ATEC as often as they wanted at their own discretion. As a result, over the time span of 12 months parents completed 12,912 ATEC evaluations.

During the data analysis stage, each ATEC was evaluated for internal consistency. Inconsistent ATEC evaluations (1,169 out of 12,912, or 9.1%) were likely the result of users who wanted access to the app but did not want to spend time to carefully complete 77 questions.

We assessed the consistency of each evaluation by looking for irregular patterns of subscale scores. The four subscales of ATEC reverse their questions in contiguous subscales so that a person with severe autism, for example, will likely answer “A. not true” to most of the questions in subscale 1, “C. very descriptive” in subscale 2, “A. not descriptive” in subscale 3, and “D. serious problem” in subscale 4. An insincere responder rushing through the questions may choose to always select the answers displayed either on the top or on the bottom of the list resulting in subscale I to subscale IV scores of 28, 0, 36, 0 or 0, 40, 0, 75, respectively. In either case, the inconsistent seesaw pattern of subscale scores is easily caught by an algorithm. This algorithm, however, would not be able to identify a responder who always selects the answer in the middle of the list. These responders were identified by calculating the standard deviation in each subscale. An insincere responder who always selects the same answer in the middle of the list will result in a standard deviation of zero for that section. Inconsistent ATEC evaluations with unusually low standard deviation and/or seesaw pattern of subscale scores were deleted from further analysis.

Autism severity was assigned using the average of all ATEC scores completed by a subject and the norms shown in Table 3. The decision to use the average ATEC score instead of the initial ATEC score was based on our observation of high intra-subject variability of ATEC scores. The standard deviation of subjects’ total ATEC score was on average 14%.

**Table 3.** ATEC autism severity norms. Percentile is from https://www.autism.com/ind_atec_report

| Table 3 |  |  |  |  |
| --- | --- | --- | --- | --- |
| Autism severity group | ATEC score | Percentile | Num of subjects in our sample | % subjects in each severity group |
| Mild | ATEC≤50 | low third: 0 to 29% | 187 | 12 |
| Moderate | 50<ATEC<80 | mid third: 30 to 70% | 675 | 46 |
| Severe | ATEC≥80 | high third: 70 to 99% | 652 | 42 |

**Table 4.** The number of subjects (and the percent of the total) in each age and severity category

| Table 4 |  |  |  |  |
| --- | --- | --- | --- | --- |
| Age group | Severity group: mild | Severity group: moderate | Severity group: severe | Totals |
| 2 to 3 years old | 101 (7%) | 343 (23%) | 343 (23%) | 787 (52%) |
| 4 to 5 years old | 51 (3%) | 217 (14%) | 185 (12%) | 453 (30%) |
| 6 to 12 years old | 35 (2%) | 115 (8%) | 124 (8%) | 274 (18%) |
| Totals | 187 (12%) | 675 (45%) | 652 (43%) | 1514 (100%) |

### Statistical Analysis

Normally distributed data were expressed as mean ± standard deviation. Non-parametric data were presented as median with interquartile ranges (IRQ). Comparisons of normally distributed data were undertaken with unpaired Student’s t-tests. A two-sided p < 0.05 was regarded as statistically significant. Comparisons of non-parametric data were calculated with the Mood’s Median Test.

## Results and Discussion

In this manuscript we report data from a feasibility study of parent-administered tablet-assisted therapy for children with ASD. We were especially interested in comparing the performance of subjects of different ages and varying ASD severities. We have measured and analyzed the performance of 1,514 children who have been working with the *Mental Imagery Therapy for Autism* (MITA) application for four to twelve months, between February of 2016 and February of 2017. We evaluated the application along four parameters: usability, feasibility, fidelity of implementation, and promise of outcomes.

### 1. Usability: Are Children Able To Understand And Physically Use The Application? Does Usability Vary Between The Different Age Groups And ASD Severity Groups?

When assessing the usability of MITA, we were particularly interested in evaluating the performance of the most vulnerable population: the newly-diagnosed 2-year-old children who may not yet be receiving adequate therapy and those with the most severe symptoms, all of whom have the most to benefit from early and intensive therapy. To assess MITA usability quantitatively we calculated the following daily averages by age and severity group: 1) the number of puzzles solved (Figure 2 and Table 5) and 2) the amount of time spent engaging with the MITA app (Figure 3 and Table 6).

**Figure 2.**
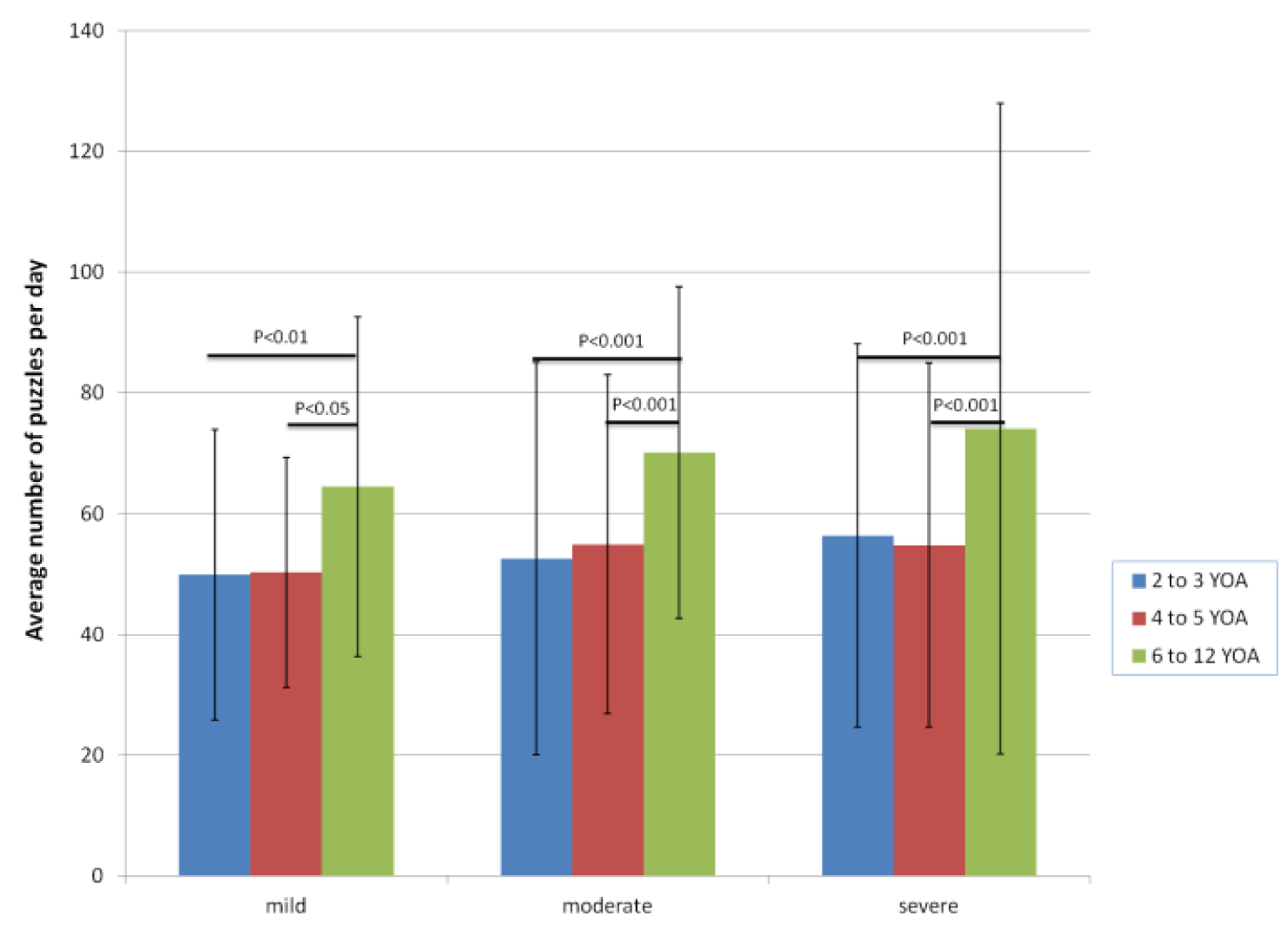
The average number of puzzles per day across the nine age and severity groups. The error bars show standard deviation. T-test was used to compare groups with the same age and the same severity. Group pairs with statistically significant difference are indicated with a horizontal bar and a corresponding P-value. There was no statistically significant difference between other groups.

**Figure 3.**
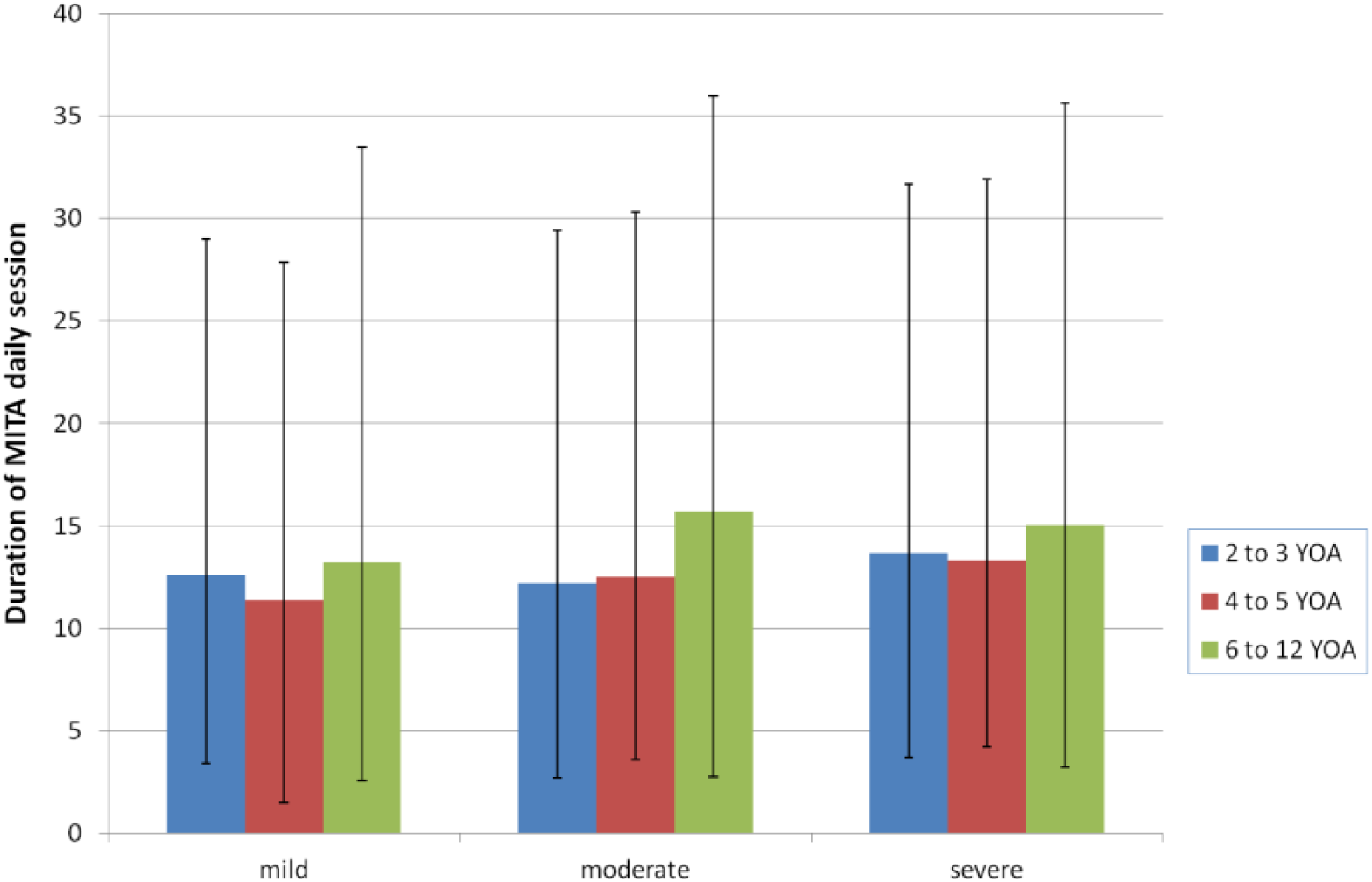
The median duration of MITA exercises per day in minutes for the nine age and severity groups. The error bars show quartile 1 and quartile 3. There was no statistically significant difference between any of the groups.

**Table 5.** Average ± SD number of puzzles per day by age and severity group

| Age group | Severity group: mild | Severity group: moderate | Severity group: severe | Totals |
| --- | --- | --- | --- | --- |
| 2 to 3 years old | 50±24 | 53±33 | 56±32 | 54±31 |
| 4 to 5 years old | 50±19 | 55±28 | 55±30 | 54±28 |
| 6 to 12 years old | 64±28 | 70±27 | 74±54 | 71±42 |
| Totals | 53±24 | 56±31 | 59±37 | 57±33 |

**Table 6.** The median (IRQ) duration of MITA exercises per day for the nine age and severity groups

| Age group | Severity group:<br>mild | Severity group:<br>moderate | Severity group:<br>severe | Totals |
| --- | --- | --- | --- | --- |
| 2 to 3 years old | 12.6 (9.2-16.4) | 12.2 (9.5-17.2) | 13.7 (10-18) | 12.9 (9.6-17.3) |
| 4 to 5 years old | 11.4 (9.9-16.45) | 12.5 (8.9-17.8) | 13.3 (9.1-18.6) | 12.8 (9.0-17.9) |
| 6 to 12 years old | 13.2 (10.65-20.25) | 15.7 (12.95-20.25) | 15.05 (11.8-20.6) | 15.1 (12.0-20.4) |
| Totals | 12.5 (9.7-16.6) | 13.1 (9.7-18.1) | 13.7 (10.2-18.6) | 13.3 (9.9-18.2) |

#### 1) Average Number of Puzzles Per Day

As described above, for children younger than 6 years of age, the MITA daily session is automatically pre-set to five puzzles in each of four activities, for a total of 20 puzzles, but children can continue solving puzzles for as long as they want. Our data show that children between the ages of 2 and 6 solved, on average, 54±30 puzzles per day, nearly three times the initial settings, which is a good indication of children’s proclivity for solving puzzles. Furthermore, pair-wise age group comparisons within each severity category showed no significant difference in the number of daily puzzles between the 2 to 3-year-old age group (54±31 puzzles on average, Table 5) and the 4 to 5-year-old group (54±28 puzzles on average), indicating that children as young as 2 can successfully use a therapeutic application regardless of ASD severity. In addition, pair-wise severity group comparisons within each age category showed no significant difference among any of the severity groups (Fig. 2), which means that subjects with severe symptoms were solving just as many puzzles as those with mild symptoms in every age group.

For children who are 6 and older, the MITA daily session is automatically pre-set to 30 puzzles (five puzzles in each of six activities). Our data show that the average number of puzzles solved per day by 6 to 12-year-olds is 71±42 (Table 5) which is more than twice the initial settings. As a result of the higher automated app settings for the older age group, pair-wise age comparisons within each severity category showed statistical significance in the number of puzzles solved per day by the 6 to 12-year-old group compared to all other age groups.

The propensity of all subjects to continue solving puzzles well past the minimum requirement necessary to earn the reward matches the anecdotal evidence gathered from parents who have reported their child’s fondness for the application.

#### 2) Average Number of Minutes Per Day

The median (IQR) amount of time spent with the MITA exercises per day was 13.3 (9.9-18.2) minutes, Fig. 3 and Table 6, which fell above our recommendation of 10 minutes per day, with no statistically significant difference among any of the age or severity groups. The small increase in the duration of MITA exercises in older children is, once again, the result of the automated app settings which require older children to complete a greater number of puzzles before earning Playtime. Notably, 19% of MITA sessions lasted more than 20 minutes, and 6% lasted more than 30 minutes, indicating that it is possible for kids to be engaged with a therapeutic app for a considerable amount of time.

The quantitative data allows us to conclude that children as young as 2 with a variety of ASD severities are able to physically use, understand and engage with the activities contained in a therapeutic app for at least 10 minutes per day, and are able to complete and exceed the length of a recommended daily session of activities.

### 2. Feasibility: Can The Application Be Used Over An Extended Period Of Time Thus Making It Feasible As A Therapeutic Application? Is It Equally Feasible For Every Age Group And Severity Level?

Over the span of our study, in which participants were able to exercise with MITA for four to twelve months, the median (IRQ) number of days each child engaged with MITA was 20 (14-42) and the median (IRQ) total number of puzzles solved was 1,317 (703-2,646). MITA usage indicates that children were able to come back to MITA day after day over the course of months. However, MITA exercises are expected to have their greatest effect in those who exercise consistently over several years.

To gauge MITA dropout rate, we wanted to focus on subjects who were serious about working through MITA therapy and exclude those who were just playing with the free app. The first significant hurdle for the parents is the first 77-quuestion ATEC evaluation, which was administered approximately one month after the first use of MITA and after the child had solved at least 100 puzzles. Accordingly, only these subjects (about 20% of all MITA downloads) were included into dropout rate calculations. Figure 4 shows the percent of subjects remaining in the study 3 months and 6 months after filling out the first evaluation. The data indicate a significant dropout rate, with only 12% of subjects remaining 6 months after the first evaluation.

**Figure 4.**
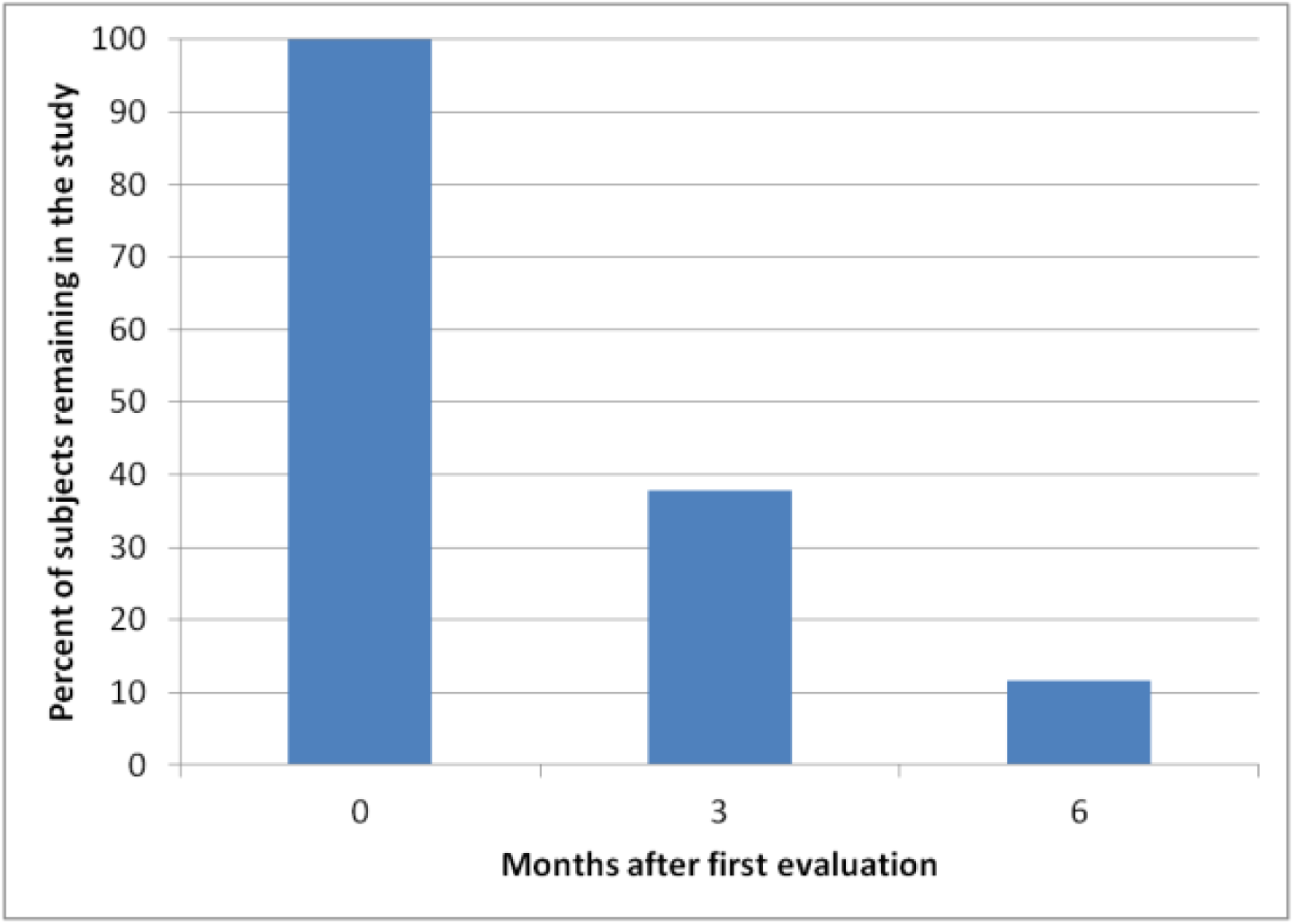
Percent of subjects remaining in the study as a function of time after the first evaluation.

We were interested in checking whether weaker subjects who showed less interest in MITA and whose parents would therefore be less motivated to fill out the questionnaire dropped out of the study at a greater rate than stronger, more engaged subjects, thus creating a positive selection process for stronger children. To address this question, we looked at the proportion of subjects who dropped out after filling out the first evaluation and before filling out the second evaluation across the nine age and severity groups, Fig. 5 and Table 7. Interestingly, among the three age groups, 2 to 3-year-olds were less likely to discontinue using MITA. Among 2 to 3-year-olds (but not among 4 to 5 and 6 to 12-year-olds), those with more severe ASD symptoms were more likely to stop using MITA. Overall, however, weaker subjects (2 to 3-year-olds with severe ASD) were dropping out at a lower rate than the strongest subjects (6 to 12-year-olds with mild symptoms). These data allow us to conclude that despite a handful of anecdotal examples of weaker subjects dropping out of the study, there is little evidence of an overall trend of positive selection of stronger children into the MITA study.

**Figure 5.**
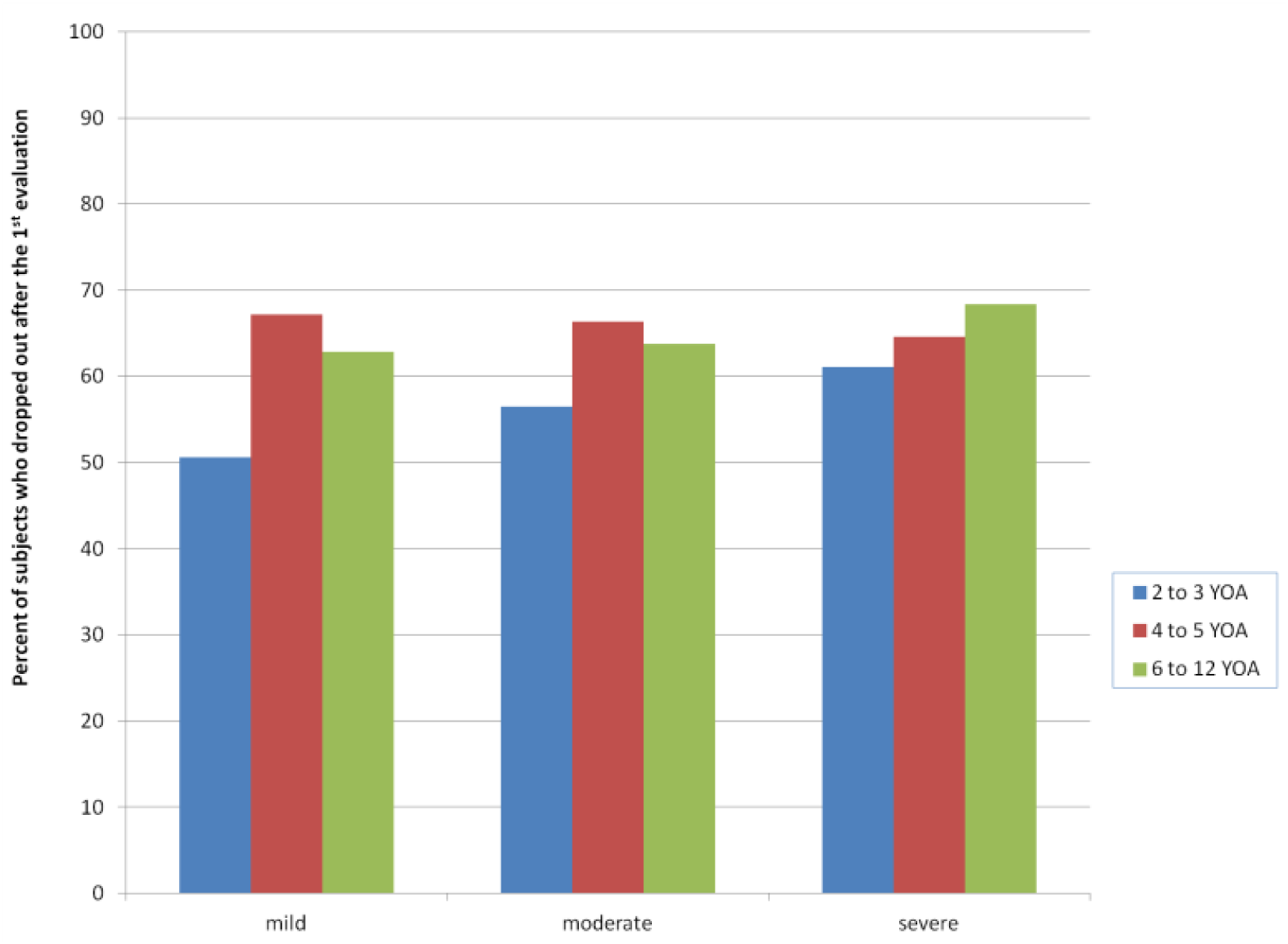
Percent of subjects who dropped out after filling out the first evaluation and before filling out the second evaluation across the nine age and severity groups.

**Figure 6.**
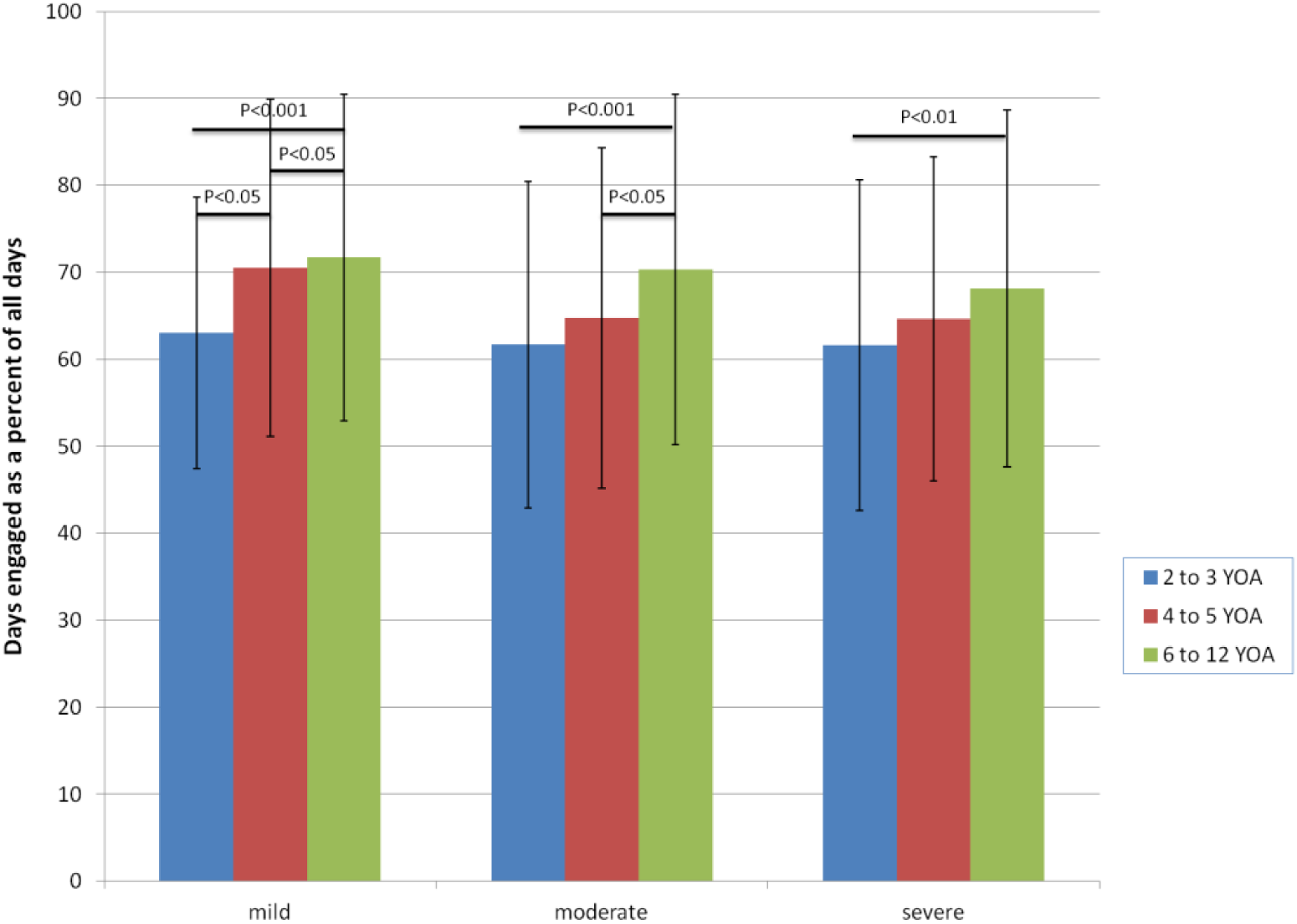
Days in which subjects completed a daily session as a percent of all days they started MITA. The error bars show standard deviation. T-test was used to compare groups with the same age and the same severity. Group pairs with statistically significant difference are indicated with a horizontal bar and a corresponding P-value. There was no statistically significant difference between other groups.

**Table 7.** Number of subjects who dropped out after filling out the first evaluation and before filling out the second evaluation across the nine age and severity groups

| Age group | Severity group:<br>mild | Severity group:<br>moderate | Severity group:<br>severe | Totals |
| --- | --- | --- | --- | --- |
| 2 to 3 years old | 86 of 170 (51%) | 352 of 623 (57%) | 490 of 802 (61%) | 928 of 1595 (58%) |
| 4 to 5 years old | 84 of 125 (67%) | 315 of 475 (66%) | 325 of 503 (65%) | 724 of 1103 (66%) |
| 6 to 12 years old | 61 of 97 (63%) | 162 of 254 (64%) | 214 of 313 (68%) | 437 of 664 (66%) |
| Totals | 231 of 392 (59%) | 829 of 1352 (61%) | 1029 of 1618 (64%) | 2089 of 3362 (62%) |

One of the main reasons cited by parents for dropping out of the study is that their child lost interest in Playtime, MITA’s primary reward structure. We are currently working to improve our incentive/reward structure by making it more varied and time-limited as well as allowing parents to personalize it to reflect their child’s unique interests, with the goal of maintaining engagement for a variety of kids over the course of years.

MITA usage data indicates it is feasible for a therapeutic application to be used over the course of months by kids as young as 2 years of age with any ASD severity, but there is likely to be significant dropout over time. For now, we have insufficient data to gauge the feasibility of multi-year usage. We are continuing our study and will be able to report in the near future.

### 3. Fidelity Of Implementation: Is The Application Being Used As It Was Intended? How Does Actual Usage Compare To Recommended Usage? Is The App Being Used Differently By Different Age Groups And ASD Severity Groups?

To assess the fidelity of implementation we compared usage recommendations with actual practice data. Our two chief recommendations were: 1) that a MITA daily session should consist of at least 20 puzzles, and that 2) children should engage with MITA on a daily basis.

1) Our recommended session length of 20 puzzles was implicit in our pre-set minimum settings. However, as we have mentioned parents have the ability to override the recommendation to as little as 5 puzzles per day, and in addition kids could quit MITA at any time. Our data show that subjects across all age and severity groups adhered to the recommendation at least 64±19% of the time, Table 8. Kids older than 6 were able to adhere to recommendations significantly better than 2 to 3-year-olds in every severity group, but this is again likely the result of the automated app settings which present older children with a greater number of puzzles before earning Playtime. Most notably, there was no significant difference in adherence to recommended session length among any of the severity groups.

2) To encourage the maximum impact of exercises, we recommended that MITA be used daily. Although there was one subjects who worked with MITA every day for over ten months, confirming that it *is* possible to engage with MITA on a daily basis, the actual median (IQR) number of days children engaged with MITA per week was 0.8 (0.5-1.5), significantly less than our recommendation. Specifically, only 200 subjects (13%) were engaged with MITA more than twice per week on average, only 72 subjects (5%) were engaged with MITA more than 3 times per week; only 28 subjects (2%) were engaged with MITA more than 4 times per week; and only 7 subjects (0.5%) were engaged with MITA more than 5 times per week. Despite these low numbers, it is interesting to note that pair-wise age and severity group comparisons showed no significant difference between the amount of usage in any of the groups, Fig. 7 and Table 9, indicating that age and severity of symptoms does not influence implementation.

**Figure 7.**
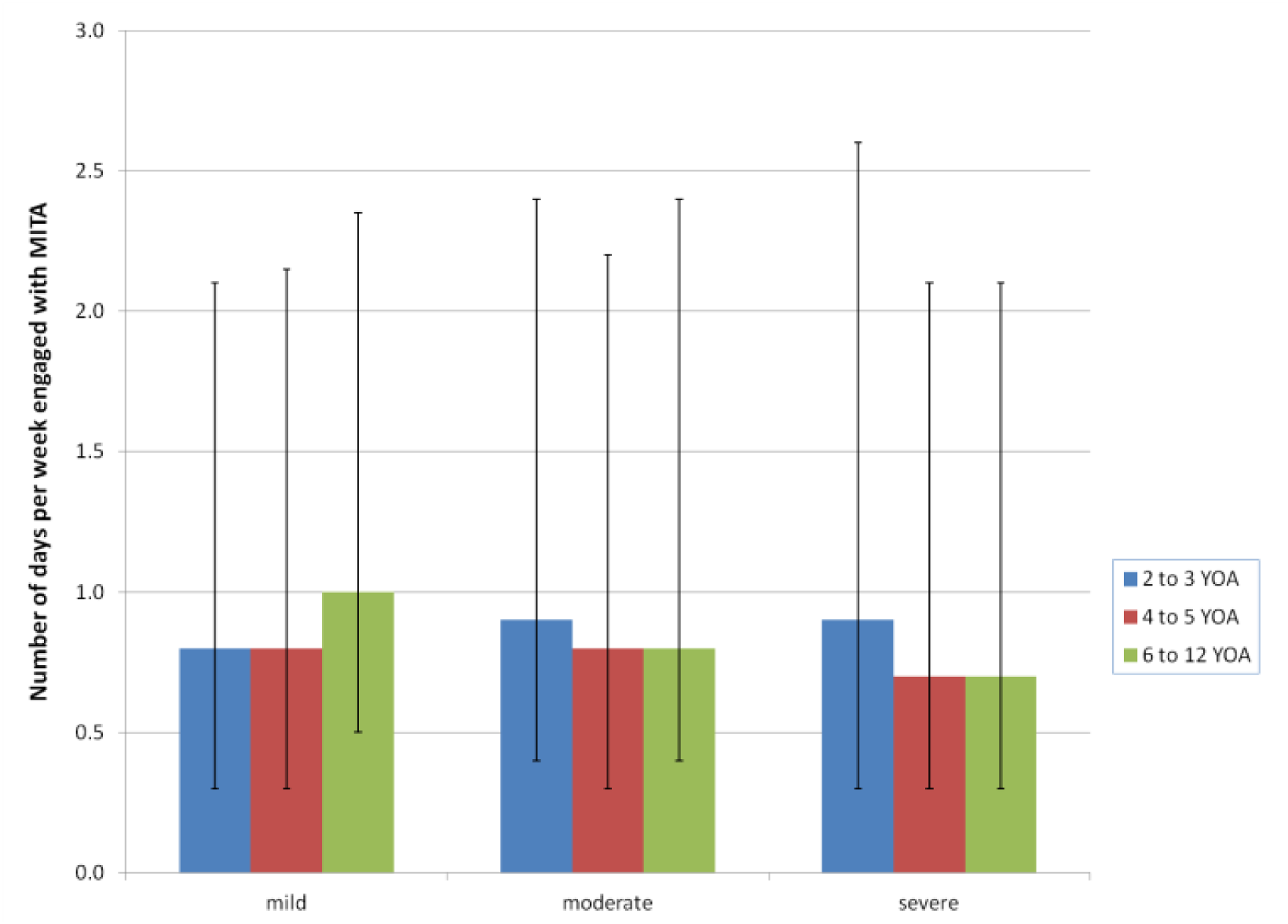
The median number of days per week engaged with MITA exercises for the nine age and severity groups. The error bars show quartile 1 and quartile 3. There was no statistically significant difference between groups.

**Table 8.** Days in which subjects completed a daily session as a percent of all days they started MITA

| Age group | Severity group:<br>mild | Severity group:<br>moderate | Severity group:<br>severe | Totals |
| --- | --- | --- | --- | --- |
| 2 to 3 years old | 63±16 | 62±19 | 62±19 | 62±19 |
| 4 to 5 years old | 71±19 | 65±20 | 65±19 | 65±19 |
| 6 to 12 years old | 72±19 | 70±20 | 68±21 | 70±20 |
| Totals | 67±18 | 64±20 | 64±19 | 64±19 |

**Table 9.** The median (IRQ) number of days per week engaged with MITA exercises for the nine age and severity groups

| Age group | Severity group:<br>mild | Severity group:<br>moderate | Severity group:<br>severe | Totals |
| --- | --- | --- | --- | --- |
| 2 to 3 years old | 0.8 (0.5-1.3) | 0.9 (0.5-1.5) | 0.9 (0.6-1.7) | 0.9 (0.5-1.6) |
| 4 to 5 years old | 0.8 (0.5-1.35) | 0.8 (0.5-1.4) | 0.7 (0.4-1.4) | 0.8 (0.5-1.4) |
| 6 to 12 years old | 1 (0.5-1.35) | 0.8 (0.4-1.6) | 0.7 (0.4-1.4) | 0.8 (0.4-1.5) |
| Totals | 0.8 (0.5-1.3) | 0.9 (0.5-1.5) | 0.8 (0.5-1.6) | 0.8 (0.5-1.5) |

Even more telling, among the subjects who engaged with MITA at least twice per week, the highest proportion was among 2 to 3 year-olds with severe ASD, Fig. 8 and Table 10. A likely explanation for this may be parents’ motivation to work with children who do not yet receive sufficient therapy. Since the vast majority of participants were from the United States where a plethora of services are available to older children diagnosed with ASD, it is possible that this pattern of use may be explained by enrollment in other therapies. Since there are fewer services available for younger children (Hume et al., 2005) parents may be more likely to fill-in the gap by working with kids on their own. When services do become available, parents may be less motivated to administer MITA on a regular basis.

**Figure 8.**
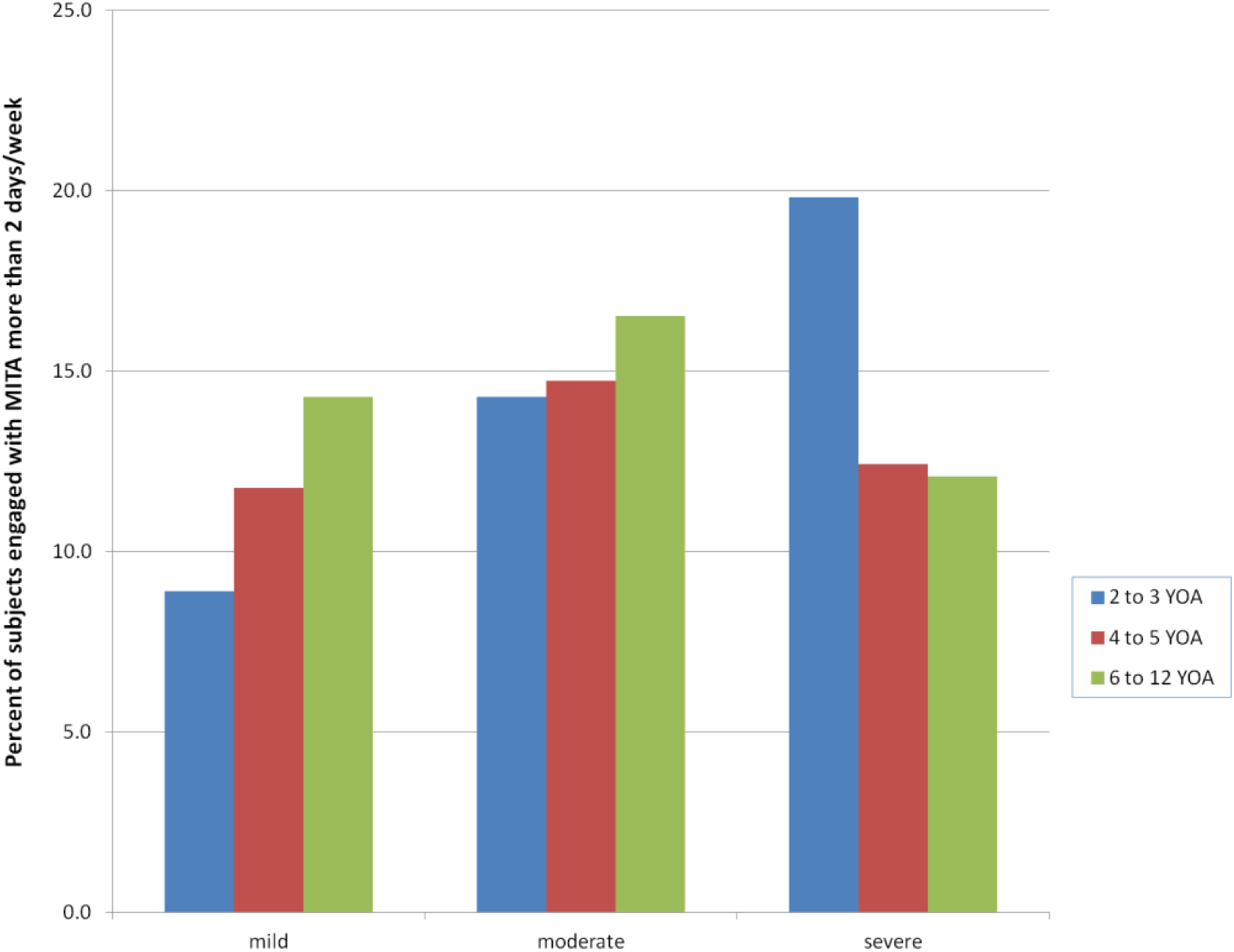
Subjects engaged with MITA more than twice per week as a percent of all subjects in that group.

**Table 10.** Subjects engaged with MITA more than twice per week as a percent of all subjects in that group

| Table 10 |  |  |  |  |
| --- | --- | --- | --- | --- |
| Age group | Severity group:<br>mild | Severity group:<br>moderate | Severity group:<br>severe | Totals |
| 2 to 3 years old | 8.9 | 14.3 | 19.8 | 16.0 |
| 4 to 5 years old | 11.8 | 14.7 | 12.4 | 13.5 |
| 6 to 12 years old | 14.3 | 16.5 | 12.1 | 14.2 |
| Totals | 10.7 | 14.8 | 16.3 | 14.9 |

The quantitative data indicates that while the application is being used as it was intended as far as the daily session length, with more than 64% of daily sessions adhering to our recommendation across all ages and severity levels, it is not being used on a daily basis as recommended. Encouragingly, however, our youngest subjects with the most severe ASD symptoms constitute the highest proportion of MITA’s “most frequent users.” It will be interesting to see if improvements in MITA’s reward structure will encourage children to engage with the application more frequently.

### 4. Promise Of Outcomes: Can A Child Show Improvement Within An Activity? Does The Extent Of Improvement Vary Between The Different Age Groups And ASD Severity Groups?

A child’s ability to improve was assessed by the Highest Achievement score and the Average Performance score, which measure a child’s best performance and average performance, respectively. In addition, we looked at and compared the number of errors per puzzle made by all nine age and severity groups as well as the daily change in performance score.

#### 1) Highest Achievement score

As described in methods, the Highest Achievement score is a measure of a child’s best performance in MITA and is equal to the sum of sustained maximum levels reached by the child in each of the nine activities normalized by the maximum possible level in each activity. The Highest Achievement score, therefore, can be thought of as a child’s “MITA IQ” as it is a gauge of overall abilities. Since subjects worked with MITA for variable duration (from 4 to 12 months) and at variable intensity (from 0.1 to 7 days a week), making a direct comparison of the Highest Achievement score is not appropriate. It is clear, for example, that reaching a Highest Achievement score of 50 in 10 days is qualitatively different than reaching the same score in 300 days. Consequently, we present the Highest Achievement score as a function of days engaged with MITA.

Each red square in the nine graphs in Figure 8 represent one subject’s Highest Achievement score plotted versus the number of days the child was engaged with MITA. Since this study is ongoing, the data in Figure 9 is a snapshot of achievement as of February 2017; most of the individuals have continued to climb up the achievement ladder. Thus, Highest Achievement score at an arbitrary date is not as telling as the speed at which a child reached that score.

**Figure 9.**
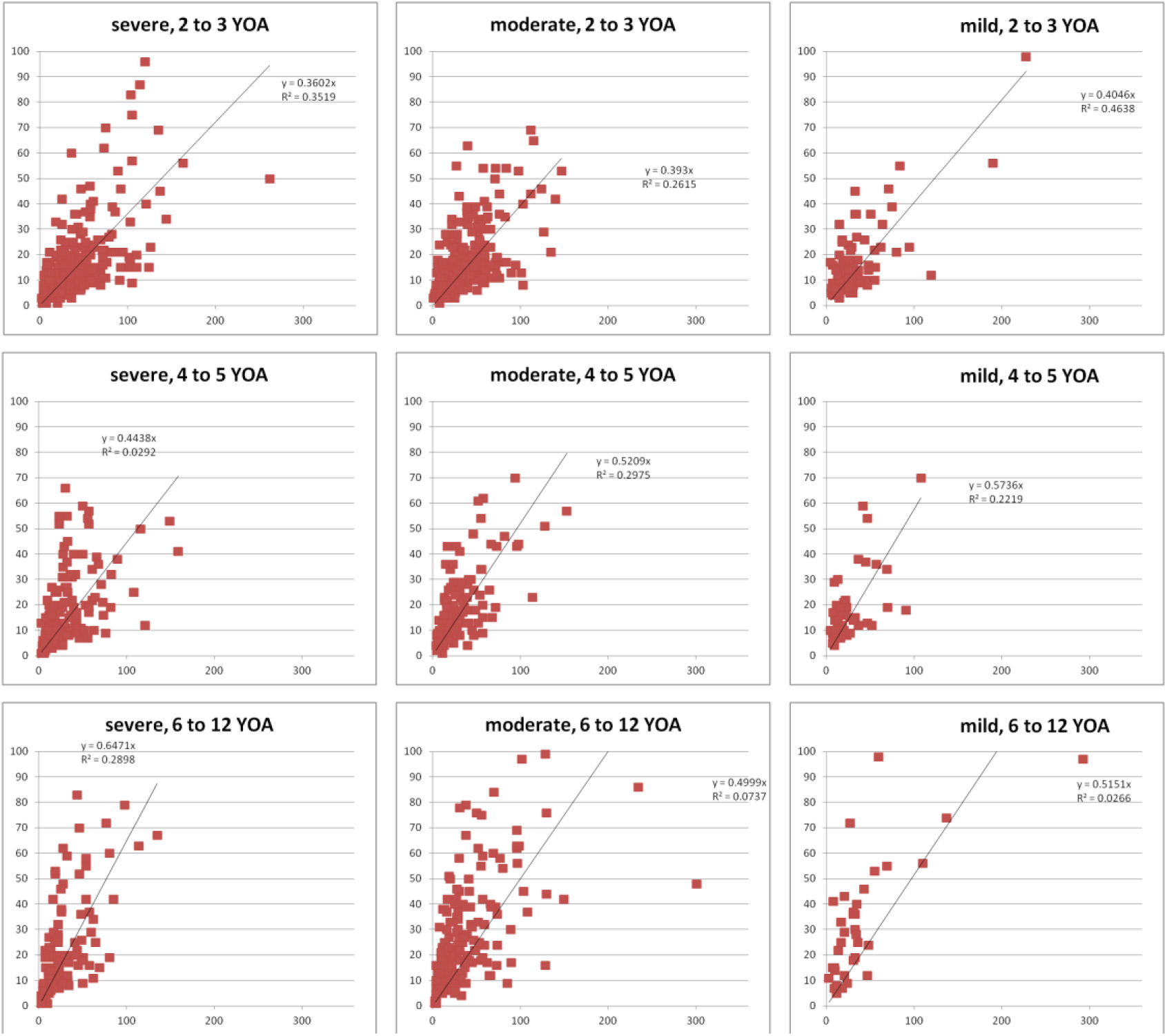
Highest Achievement score vs. number of days engaged with MITA

Figure 10 and Table 11 compare the speed of achievement (average Highest Achievement score increase per daily session) for all nine age and severity groups. All age groups and severity levels were able to increase their Highest Achievement score over time, but 6 to 12-year-olds and 4 to 5-year-olds increased their score at a significantly faster rate than 2 to 3-year-olds (across all severities) and faster than 4 to 5-year-olds in the moderate and severe categories. In other words, older children working with MITA climbed along MITA difficulty levels quicker than younger children, regardless of ASD severity. This result is not surprising, since we would generally expect older children to have higher overall achievement speed. However, we were surprised to see that there was no significant difference in the speed of achievement within any of the age groups. In other words, 2 to 3-year-olds had similar speeds of achievement regardless of whether their symptoms are mild, moderate or severe; the same is true for every other age group in our study. This data is very encouraging as it suggests that MITA exercises are just as accessible and beneficial to the most needy population (individuals with severe symptoms) as it is to those with milder symptoms.

**Figure 10.**
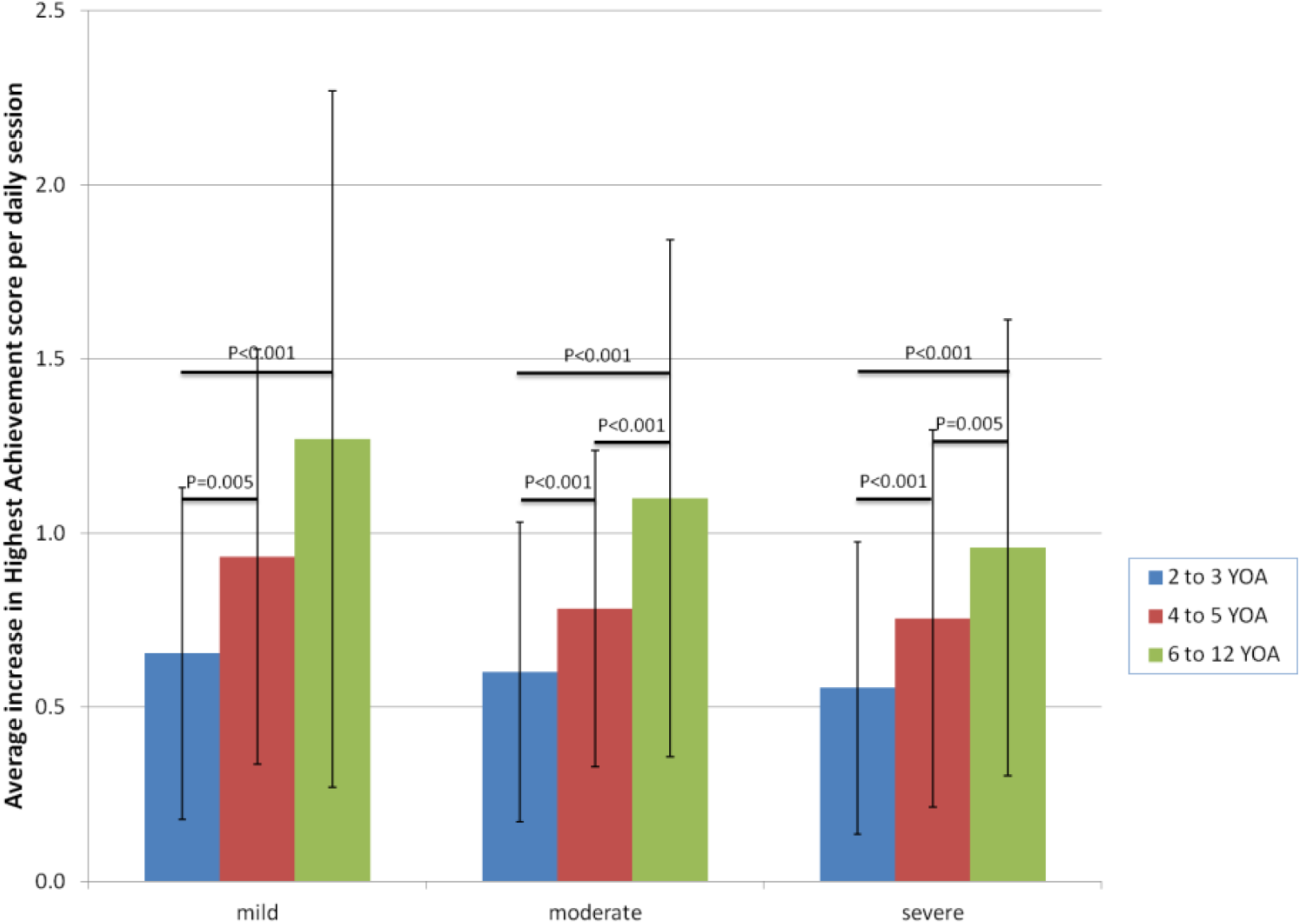
The speed of achievement, calculated as the average increase in Highest Achievement score per daily session. A speed of achievement of 1 approximately corresponds to the cumulative increment of six levels per day across all activities. The error bars show standard deviation. T-test was used to compare groups with the same age and the same severity. Group pairs with statistically significant difference are indicated with a horizontal bar and a corresponding P-value. There was no statistically significant difference between other groups.

**Table 11.**
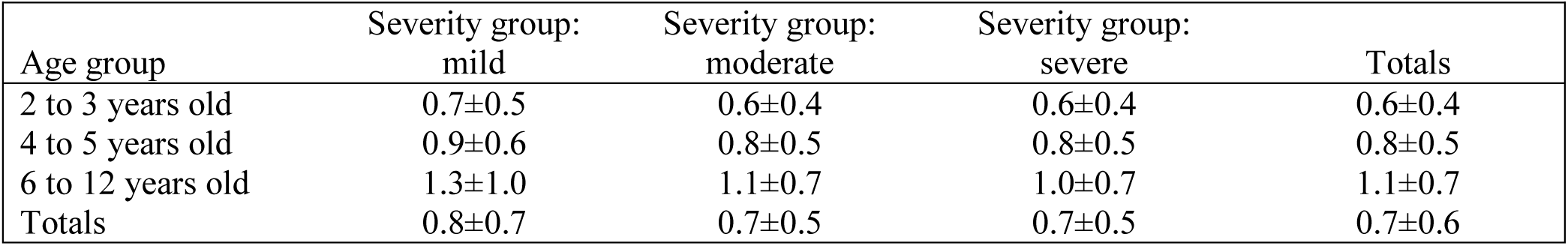
The speed of achievement, calculated as the average increase in Highest Achievement score per daily session

#### 2) Average Performance score

Unlike the Highest Achievement score which considers a child’s best performance, the Average Performance score is a measure of overall performance over all daily sessions. The Average Performance score, therefore, can be thought of as a child’s cumulative “MITA grade.” As educators, we expected to see a normal Gaussian distribution with an average in the C range (70-80%), and we were hoping that none of our “students” were failing the MITA course. The Average Performance score data, Fig. 11 and Table 12, met our expectations. All nine age and severity groups had average scores between 70 and 78 but 6 to 12- year-olds with mild symptoms did significantly better than all comparable groups and older kids generally did better within each severity group. However, just as with the Highest Achievement score, we were surprised that there were no significant differences within the 2 to 3- and 4 to 5- year-old age groups, meaning that ASD severity did not affect the overall performance scores.

**Figure 11.**
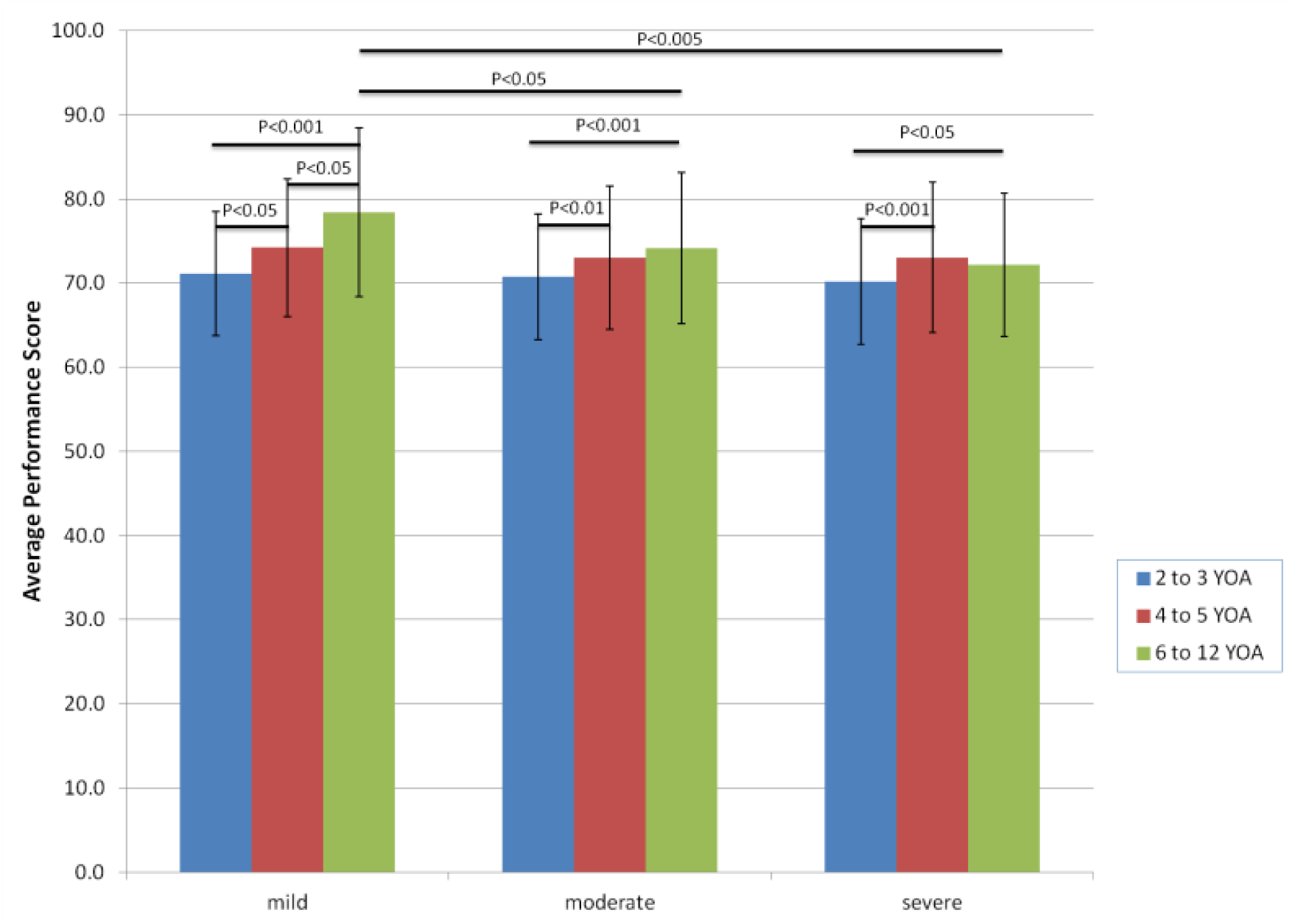
The Average Performance score across the nine age and severity groups. The error bars show standard deviation. T-test was used to compare groups with the same age and the same severity. Group pairs with statistically significant difference are indicated with a horizontal bar and a corresponding P-value. There was no statistically significant difference between other groups.

**Table 12.**
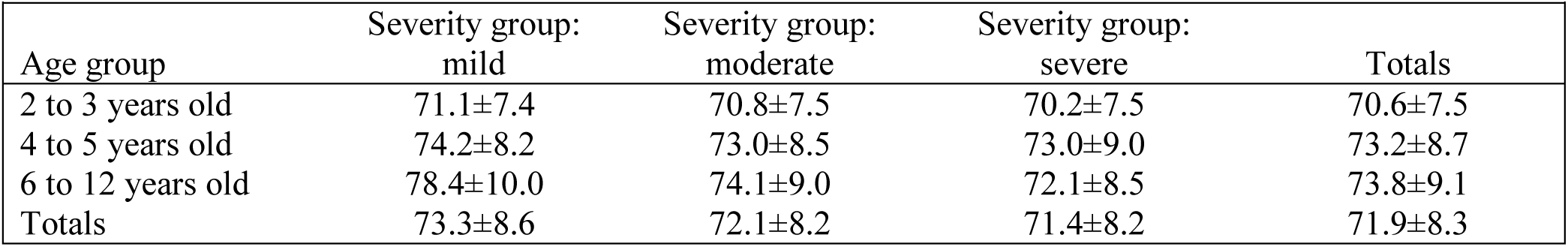
The Average Performance score across the nine age and severity groups

| Age group | Severity group:<br>mild | Severity group:<br>moderate | Severity group:<br>severe | Totals |
| --- | --- | --- | --- | --- |
| 2 to 3 years old | 71.1±7.4 | 70.8±7.5 | 70.2±7.5 | 70.6±7.5 |
| 4 to 5 years old | 74.2±8.2 | 73.0±8.5 | 73.0±9.0 | 73.2±8.7 |
| 6 to 12 years old | 78.4±10.0 | 74.1±9.0 | 72.1±8.5 | 73.8±9.1 |
| Totals | 73.3±8.6 | 72.1±8.2 | 71.4±8.2 | 71.9±8.3 |

#### 3) Number of Errors per Puzzle

While the Average Performance Score accounts for the number of errors made per puzzle normalized by the number of answer choices, we wanted to look directly at error rate as an additional measure of performance. The quantitative data shows that, in general, the youngest subjects make a significantly greater number of mistakes than older subjects, Fig. 12 and Table 13. Within the 6 to 12-year-old age group, kids with more severe symptoms also made more mistakes. However, just as with the other performance data, we were surprised and encouraged that there was no significant differences within the 2 to 3- and the 4 to 5-year-old age groups, meaning that the severity of symptoms did not affect the error rate.

**Figure 12.**
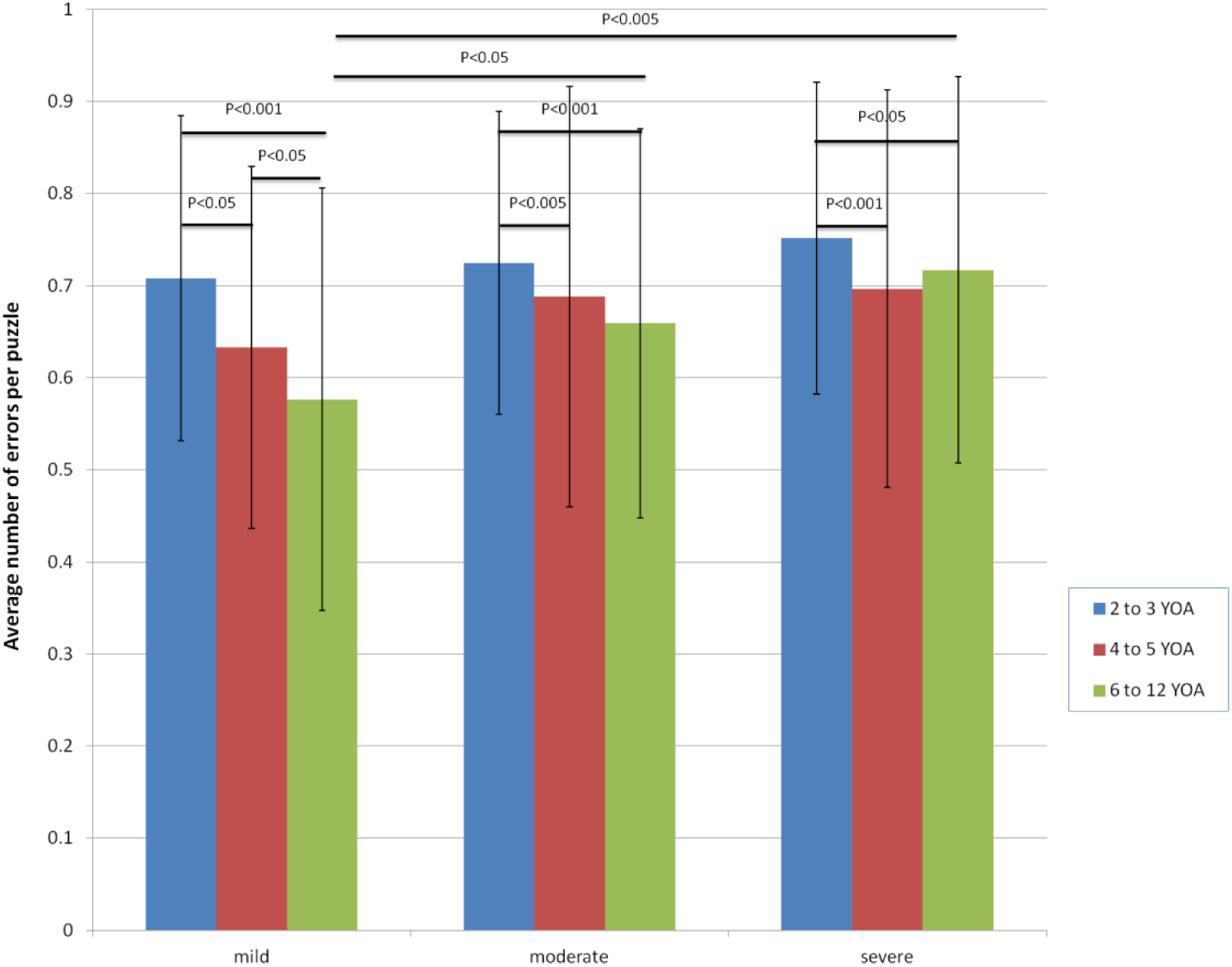
The average number of errors per puzzle across the nine age and severity groups. The error bars show standard deviation. T-test was used to compare groups with the same age and the same severity. Group pairs with statistically significant difference are indicated with a horizontal bar and a corresponding P-value. There was no statistically significant difference between other groups.

**Table 13.** The Average number of errors per puzzle across the nine age and severity groups

| Age group | Severity group:<br>mild | Severity group:<br>moderate | Severity group:<br>severe | Totals |
| --- | --- | --- | --- | --- |
| 2 to 3 years old | 0.71±0.18 | 0.72±0.16 | 0.75±0.17 | 0.73±0.16 |
| 4 to 5 years old | 0.63±0.20 | 0.69±0.23 | 0.70±0.22 | 0.69±0.22 |
| 6 to 12 years old | 0.58±0.23 | 0.66±0.21 | 0.72±0.21 | 0.68±0.22 |
| Totals | 0.66±0.20 | 0.70±0.20 | 0.73±0.19 | 0.71±0.20 |

Arguably the most important parameter to assess the promise of outcome is the change of the Average Performance score per daily session. On average, subjects improved their Average Performance score by 0.15±0.58 per day, which translates into an impressive improvement of 4.5% over 30 days of MITA use. All groups but the oldest children with severe ASD showed some improvement in their Average Performance score. Younger subjects showed significantly more improvement than 6 to 12 year-olds in the moderate and severe categories.

Though it is true that older kids were generally doing more difficult puzzles, it is important to note that the Average Performance score is a normalized score that should not be influenced by puzzle difficulty. To confirm this, we plotted the Average Performance score as a function of puzzle difficulty for each MITA activity (data not shown). We found no correlation between the score and puzzle difficulty confirming optimal normalization. Thus, we can conclude that the improvement of the Average Performance score over time represents a genuine improvement in performance.

## Conclusions

In this manuscript we report data from a feasibility study of parent-administered tablet-assisted therapy for children with ASD. To our knowledge, it is the first study of tablet-based cognitive exercises intended as an early intervention for subjects with ASD as young as 2 years of age, and the largest of its kind (compare to Ref.(Venkatesh, Phung, Duong, Greenhill, & Adams, 2013). By looking at the performance of 1,514 children who worked with the *Mental Imagery Therapy for Autism* (MITA) computerized learning application for four to twelve months, we conclude that 1) MITA works as designed; 2) Parents are capable of administering tablet-based therapy; 3) children with ASD were able to engage with the MITA application independent of ASD severity; 4) children as young as 2 years of age (as well as older children) were able to engage with MITA exercises.

Surprisingly, the effects of ASD severity on performance were minor: Children of all ASD severity levels:

1. were climbing along difficulty levels at a comparable rate (Fig. 10 and Table 11, no statistically significant difference),
2. had comparable Average Performance scores (Fig. 11 and Table 12, statistically significant difference only for 6 to 12-year-olds),
3. had comparable number of errors per puzzle (Fig. 12 and Table 13, statistically significant difference only for 6 to 12-year-olds), and
4. had comparable daily improvements in their Average Performance score (Fig. 13 and Table 14, no statistically significant difference).

**Figure 13.**
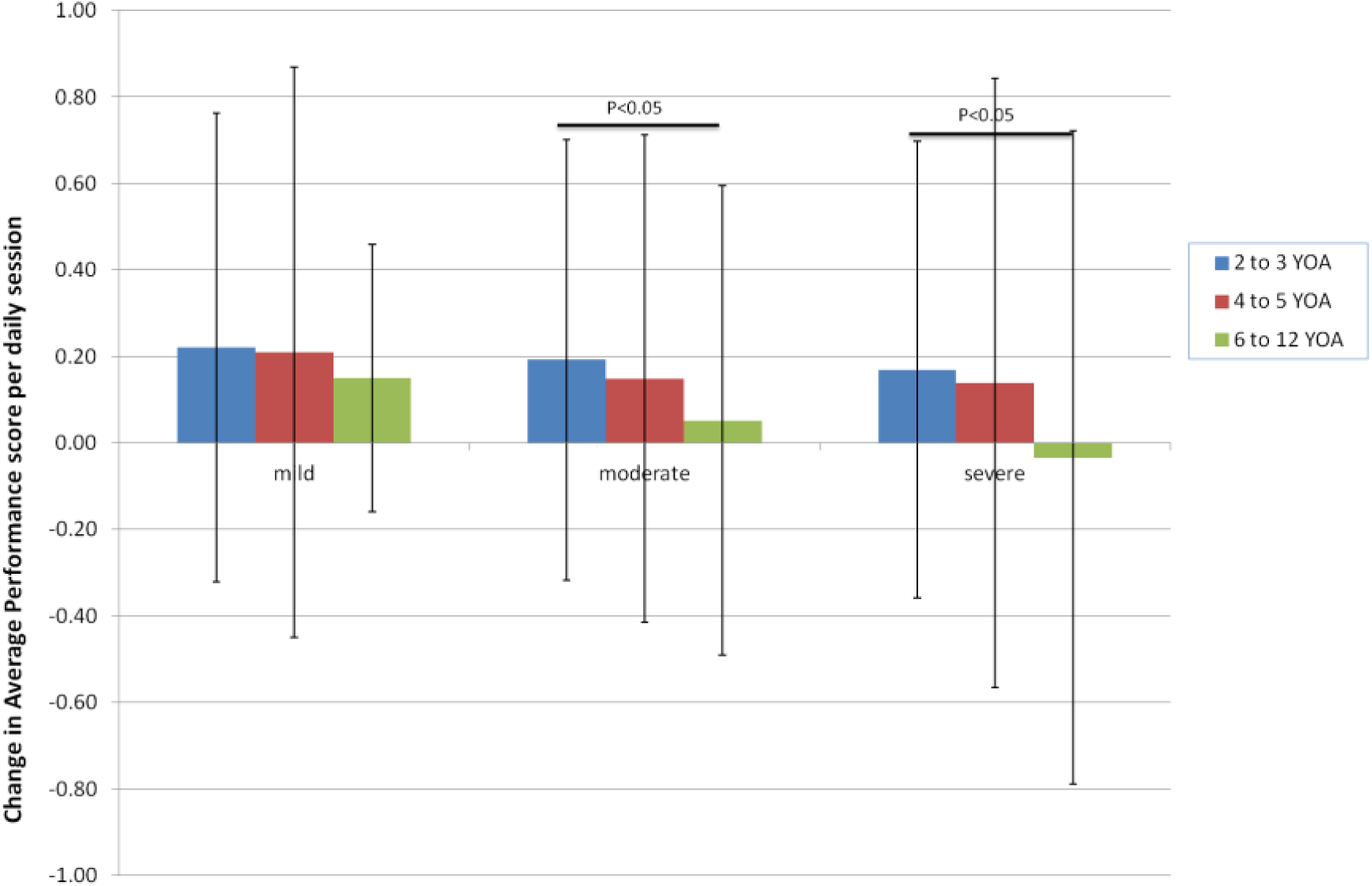
The change in the Average Performance score per daily session for all nine age and severity groups. The error bars show standard deviation. T-test was used to compare groups with the same age and the same severity. Group pairs with statistically significant difference are indicated with a horizontal bar and a corresponding P-value. There was no statistically significant difference between other groups.

**Table 13.** The change in the Average Performance score per daily session for all nine age and severity groups

| Age group | Severity group:<br>mild | Severity group:<br>moderate | Severity group:<br>severe | Totals |
| --- | --- | --- | --- | --- |
| 2 to 3 years old | 0.22±0.54 | 0.19±0.51 | 0.17±0.53 | 0.19±0.52 |
| 4 to 5 years old | 0.21±0.66 | 0.15±0.56 | 0.14±0.70 | 0.15±0.63 |
| 6 to 12 years old | 0.15±0.31 | 0.05±0.54 | -0.03±0.75 | 0.03±0.63 |
| Totals | 0.20±0.54 | 0.16±0.53 | 0.12±0.63 | 0.15±0.58 |

However, we were very encouraged by data which indicate that the severity of symptoms did not influence the performance in younger age groups. In other words, 2 to 3- and 4 to 5-year-olds showed promise of outcomes regardless of whether their ASD symptoms were mild, moderate or severe. This suggests that MITA exercises are just as accessible and beneficial to the most needy population (individuals with severe symptoms) as it is to those with milder symptoms, as long as they start exercising at a young age.

In general, the effect of age on performance was much stronger than the effect of ASD severity. Younger children:

1. showed a significantly slower increase of the Highest Achievement score (Fig. 10 and Table 11);
2. had significantly worse Average Performance scores (Fig. 11 and Table 12); and
3. had a significantly greater number of errors per puzzle (Fig. 12 and Table 13). However younger kids also showed a significantly greater improvement in the Average Performance score each day they were engaged with MITA (Fig. 13 and Table 14), underlying a considerable promise of outcomes for younger children.

It is also noteworthy that children with more severe ASD solved slightly more puzzles per day (Fig. 2 and Table 5, no statistically significant difference) and spend slightly more time with MITA per day (Fig. 3 and Table 6, no statistically significant difference), which is likely an indication of greater parental motivation in administering therapeutic exercises to this population.

The quantitative data gathered in this study allows us to make the following conclusions about usability, feasibility, fidelity of implementation and promise of outcomes:

1. **Usability:** Children as young as 2 with any ASD severity are able to physically use, understand and engage with the activities contained in the MITA therapeutic app for at least 10 minutes per day, and are able to complete more than 50 cognitive exercises presented in puzzle form.
2. **Feasibility:** It is feasible for a therapeutic application to be used over the course of months by kids as young as 2 years of age with any ASD severity, but there is likely to be significant dropout over time which may be curbed by a varied and time-limited reward structure that can be personalized by parents to reflect the unique interests of their child. For now, we have insufficient data to gauge the feasibility of multi-year usage.
3. **Fidelity of Implementation:** Subjects are capable of using the application as intended once they have sat down for a daily session, but more work needs to be done to ensure that subjects are using the application on a frequent enough basis to result in a therapeutic benefit.
4. **Promise of Outcomes:** Subjects of all ages and ASD severity were able to show some improvement within MITA by progressing through MITA difficulty levels and increasing their Highest Achievement Score over time. The greatest promise of outcome, however, is for the youngest subjects, who demonstrated the greatest improvements in their Performance Score as compared to older subjects.

MITA effectiveness is currently being tested in a longitudinal observational clinical trial, the goal of which is to test the hypothesis that regular practice with MITA will result not only in a greater ability to attend to multiple cues, but also in vast improvements on transfer tasks measuring visuo-spatial as well as communicative skills. Furthermore, we hope to show that MITA, coupled with an effective vocabulary training program, can lead to improvements in the realm of language comprehension. We predict that children who begin training at an early age, and who make consistent progress over the course of training, will see drastic improvements in their language function. Since many kids diagnosed with ASD are already receiving ample vocabulary training, what’s missing is the skill to attend to various syntactic combinations of learned words, i.e. true flexible syntactic language comprehension. For example, a child who has learned the words “fish,” “ate” and “cat” but who cannot mentally synthesize those words into a novel scene, would struggle to understand the difference between “a fish ate a cat” and “a cat ate a fish”, while a similar child who has learned mental synthesis will be successful with the task. We believe that combining the mental synthesis ability with vocabulary knowledge will result in an understanding of a full syntactic language, which will eventually lead to a significant reduction of the severity of the ASD diagnosis and ultimately to a more productive and independent life.

## Compliance with Ethical Standards

This is observational study is exempted from IRB and informed consent according to Code of Federal Regulations, TITLE 45, PUBLIC WELFARE, DEPARTMENT OF HEALTH AND HUMAN SERVICES, PART 46, PROTECTION OF HUMAN SUBJECTS, Subpart A, Basic HHS Policy for Protection of Human Research Subjects, §46.101 (b) (1): “Research conducted in established or commonly accepted educational settings, involving normal educational practices, such as (i) research on regular and special education instructional strategies, or (ii) research on the effectiveness of or the comparison among instructional techniques, curricula, or classroom management methods.”

